# Augmenting Transcranial Magnetic Stimulation Coil with Magnetic Material: An Optimization Approach

**DOI:** 10.1101/2022.01.21.477303

**Authors:** Charles Lu, Zhi-De Deng, Fow-Sen Choa

## Abstract

Transcranial magnetic stimulation (TMS) is a neuromodulation technique that has been approved by the U.S. Food and Drug Administration for several neuropsychiatric disorders, including major depression and obsessive-compulsive disorder. However, the therapeutic efficacy of TMS treatment has been modest, despite decades of research. While there are many potential reasons as to why, one of the most obvious is the limitations of current technologies. One prominent example is the penetration depth–focality tradeoff of existing TMS coils. The most widely used figure-of-8 coils stimulate brain regions just superficially under the coil, missing deep brain regions known to be critically involved in psychiatric disorders; while ring-type coils can stimulate deep into the brain, but stimulate a large brain volume (lack of focality). A new coil design strategy is proposed: magnetic materials encompassing the human head are optimized to shape the electromagnetic field generated by the primary coil. Specifically, a mathematical model was developed to describe the physical problem; the magnetic materials were discretized into unit blocks; Newton’s gradient descent method was applied to iteratively optimize the spatial distribution of the unit blocks to achieve a desired electric field distribution inside a head model. Results reveal that the proposed design achieves a coil penetration depth equal to or better than state-of-the art commercial coils, while improving the depth–focality tradeoff by a factor of 2.2 to 2.7. As a proof-of-concept, a prototype coil and a spherical head model were constructed; the spatial distribution of the induced electric field inside the head model was mapped. Results validated the proposed coil design. TMS coils based on this novel design strategy could potentially lead to better therapeutic outcome.

## 1. Introduction

Mental illnesses and substance use disorders produce a significant burden on patients and society, with an estimated national cost of about 280 billion US dollars in the year 2020 alone [1]. More specifically, treatment-resistant major depression affects approximately 2% of the general population [2]. Additionally, the ongoing national opioid crisis points to limitations of current addiction treatment, calling for novel therapeutic strategies [3, 4].

Transcranial magnetic stimulation (TMS) is a non-invasive neuromodulation technique that emerges as a potential treatment option. TMS applies strong but brief electric current pulses through a coil placed near one’s head; the coil generates magnetic fields that pass through the skull and induces electric current in the brain, stimulating the neuronal tissues. The U.S. Food and Drug Administration has approved TMS for treatment-resistant major depression, obsessive-compulsive disorder, and recently nicotine addiction [5, 6, 7]. However, therapeutic efficacy of existing TMS treatment remains modest, both the response rate and remission rate are only about 10% higher than the sham control group [2, 8]. Thus, enhancing therapeutic efficacy and utility of the TMS treatment could be of great value.

While there are many potential reasons for the modest efficacy, technical limitations, in particular, could be a major factor [9]. One such limitation is the limited penetration depth of TMS coils, which is the critical element that defines the distribution of the induced electric field (E-field), and thus which brain regions are stimulated. The most widely used clinical coil, the figure-of-8 coil, consists of two ring-shape coils placed in a figure-8 formation. This type of coil stimulates brain regions just superficially under the coil [10], missing deep brain regions known to be critically involved in psychiatric disorders. Theoretically, one can stimulate deeper structures by applying stronger electrical current to the coil, but this strategy invariably stimulates a larger brain volume which risks seizure in patients, and is thus not allowed for safety considerations [11]. There have been numerous efforts to design TMS coils that penetrate deeper into the brain while minimizing stimulated brain volume (i.e. field spread) [9, 12, 13, 14]. For example, several groups proposed the use of coil arrays, but these designs remain theoretical [13, 15]. Deng et al. systematically analyzed penetration depth–field spread tradeoff of 50 coil designs [16], and found that existing TMS coil designs fall into two categories: ring-type coils can stimulate deep into the brain, but stimulate a large brain volume (lack of focality); the figure-of-8 type coils stimulate more focal area, but only the superficial brain regions can be stimulated (lack of stimulation depth). An increase of stimulation depth results in a decrease in focality, and an increase in focality results in a decrease of stimulation depth. This is known as the depth–focality tradeoff.

In the present study, we propose an augmentation strategy that incorporates magnetic materials that are used to shape the electromagnetic field generated by the TMS coil. We hypothesized that by optimizing the spatial distribution of magnetic materials surrounding the coil, one can improve the penetration depth–field spread tradeoff. A mathematical model was developed to describe the physical problem; Newton’s gradient descent method was applied to optimize the distribution of the magnetic materials. Experimental measurements were performed that validated the proposed coil design strategy.

## 2. Methods

### 2.1. Theory

#### 2.1.1. Shaping magnetic field and induced electric field with magnetic material

As illustrated in figure 1, the magnetic field **H** at point *p* produced by coils A and B, in the presence of magnetic material C, can be expressed as [17]:

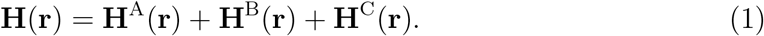

**Figure 1.**
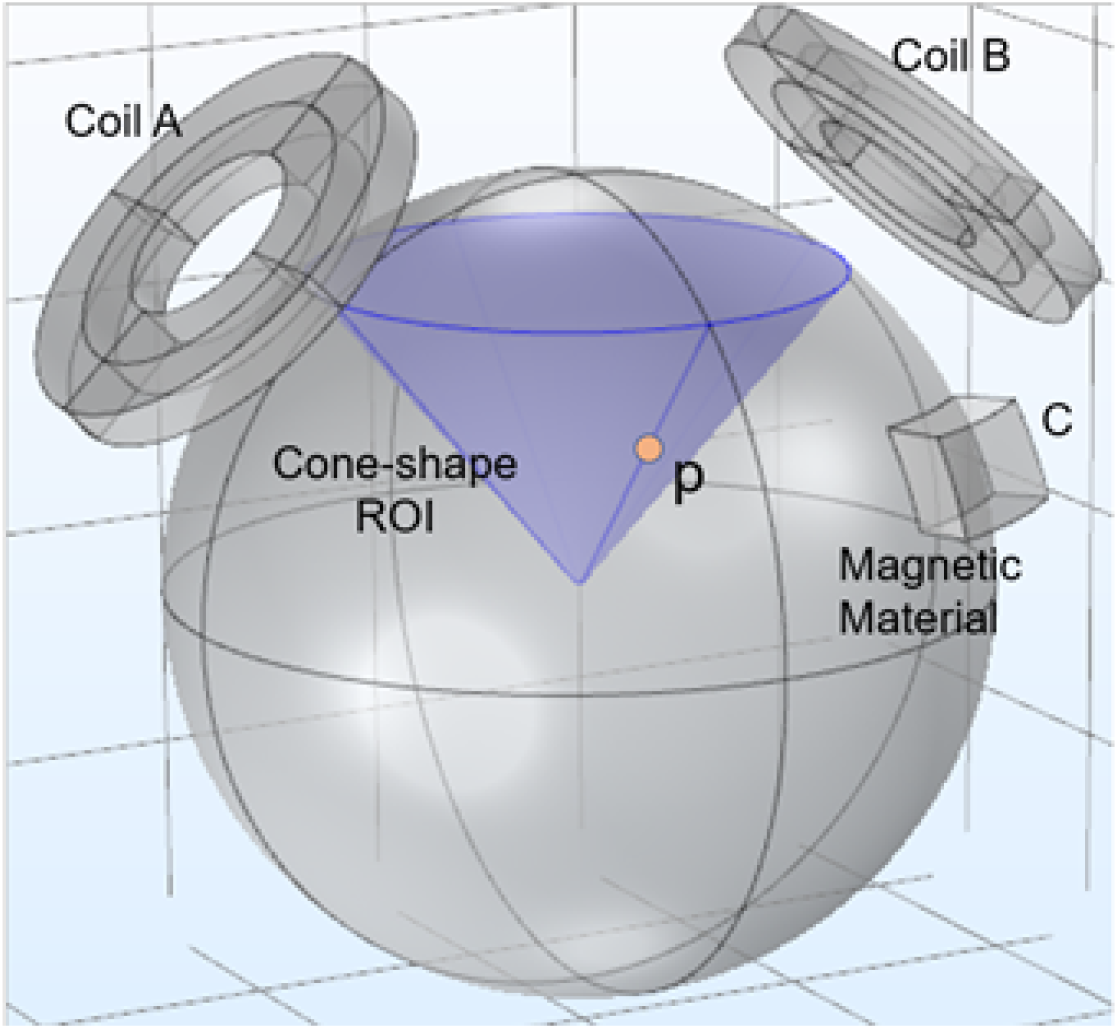
Illustration of the coil design concept. A cone-shape region-of-interest (ROI) was defined inside a spherical head model. The magnetic and electric field at any given point p in the ROI is the combined contribution from coils A, B and magnetic material C. The E-field generated by A and B can be shaped and optimized with magnetic materials to achieve desired distribution.

For coil A with current density **j**(**r′**) in a small volume element **dr′**, located at **r′**, **H**^A^(**r**) can be expressed as the following:

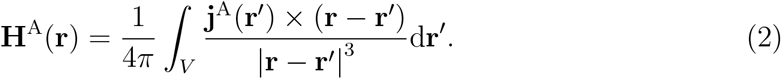

**H**^B^(**r**) has a similar expression as equation 2. Assuming coils A and B induces surface magnetic charge density *ρ*_S_(**r**) in magnetic material C, and for an isotropic magnetic material, **H**^C^(**r**) can be expressed as:

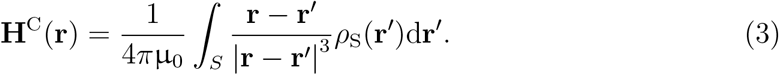

The magnetic flux **B** can be expressed as:

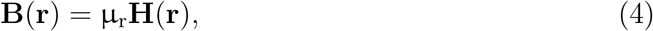

where μ_r_ is the relative permeability, depending on the properties of the magnetic of the material. It equals 1 in free space and the theoretical value for pure iron can reach 5000.

Equation 1 dictates that the magnetic field at any points *p* is the linear superposition of the field by coils and magnetic materials; equation 4 reveals that a magnetic material with a high relative permeability can effectively shape the total field generated by coils A and B. According to Faraday’s law, the induced E-field must also be altered with the addition of magnetic material C.

#### 2.1.2. Optimizing the spatial distribution of the magnetic material for optimal tradeoff in penetration depth–focality of the induced E-field

TMS-induced E-fields are always stronger on the surface than the inside of a spherically symmetric volume conductor [18]. Nevertheless, by optimizing the sizes and locations of the magnetic materials, both the decay rate, which is inversely related to penetration depth, and the field spread—which is inversely related to E-field focality—can be optimized. Specifically, the cost function was defined as:

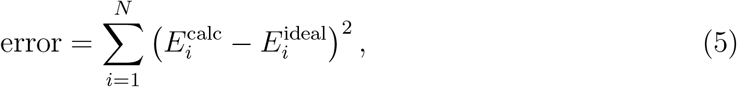

where *N* represents the total number of points in the ROI. 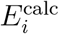 and 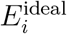 are the calculated and the desired E-field at point *i*, respectively. The goal of the optimization is to minimize the cost function.

### 2.2. E-field calculation

Since brain tissue is electrically conductive, whereas the air and skull are almost complete insulators, a time-varying magnetic field will induce an accumulation of electrical charges in tissue interfaces. The charges will generate a secondary E-field (**E**^s^), in addition to the primary E-field (**E**^p^). The total field in the brain tissue **E** is the vectorial summation of these two fields. Therefore, while the magnetic field, **H**, can be relatively straightforward to calculate, the E-field necessitates the use of specialized software.

Makarov et al. recently developed an E-field computation software package [19]. This package applies charge-based boundary element fast multipole method (BEM-FMM). The BEM-FMM approach provides unconstrained numerical field resolution close to and across material interfaces of different permittivity. The computational code was written in MATLAB, and is freely available via GitHub (https://tmscorelab.github.io/TMS-Modeling-Website/).

The human head was modeled by a sphere with a radius of 8.5 cm and isotropic conductivity of 0.33 S/m [16]. This conductivity value has been shown to be the average conductivity of human brain tissue [20]. The cortical surface was set at a depth of 1.5 cm from the surface of the head, representing the average scalp and skull thickness of an adult. The distinct head tissue layers (scalp, skull, corticospinal fluid, and brain) were not differentiated, since magnetically induced E-field in a sphere is insensitive to radial variations of conductivity [21]. The spherical head model serves as a benchmark, allowing for comparisons of coil performance among different designs, as demonstrated previously [13, 16]. The magnetic material was assumed isotropic with relative permeability 1000 and the frequency was set to 5 kHz, since it is the center frequency of a typical TMS pulse [16].

### 2.3. Quantification of penetration depth and field spread

Two parameters, *d*_1/2_ and *s*_1/2_, were used to quantify penetration depth and field spread [13, 16]. Specifically, Deng et al. [16] operationally defined *d*_1/2_ as the radial distance from the cortical surface to the deepest point where the E-field strength *E* is half of its maximum value, *E*_max_. Field spread was defined as *s*_1/2_ = *V*_1/2_/*d*_1/2_, where *V*_1/2_ is the half-value volume of the brain region that is exposed to an E-field as strong as or stronger than half of *E*_max_.

### 2.4. Coil parameters

As a starting point for the optimization, we chose to shape the field of a double-cone coil, which already has a deep field penetration. The double-cone coil configuration, a variation of the figure-of-8 coil, consisting of two ring-type coils arranged at an angle smaller than 180°, is known to have deeper penetration depth than the conventional figure-of-8 coil [14]. The inner radius of the ring coil was set to be 5 cm, outer radius 8 cm, with 6 turns of wires, and a cross sectional area of 3 × 4 mm^2^. The angle alpha between the two rings (see figure 1) was determined as follows: we modeled the E-field for each angle from 100° to 140° with a step of 5^*°*^, and calculated *s*_1/2_ and *d*_1/2_. An angle of 120° gave best tradeoff between *s*_1/2_ and *d*_1/2_ and was chosen for subsequent optimization.

### 2.5. Discretization of magnetic materials

Magnetic material that surrounded only the top half of the spherical head model was prescribed, as the lower half is too far away from the coil, and is expected to have negligible effects on the E-field (see figure 2). The magnetic material was discretized as follows: the polar angle (*θ*) of the top half was evenly divided into 6 segments, called layers. The *θ* values for layers 1 to 6 were: 0°–15°, 15°–30°, 30°–45°, 45°–60°, 60°–75°, and 75°–90°. Each layer was evenly divided along the azimuthal angle *ϕ*. Considering that the sizes of these 6 layers were quite different, *ϕ* was arbitrarily defined into different sections across these layers to achieve a similar volume for each block, with a total of 64 symmetrically distributed in the *X*–*Y* plane. Each block had an inner radius *r* of 9 cm (leaving 5 mm space between the magnetic material and the spherical head model); the thickness of each block was initially set to 1.5 cm (Δ*r*).

**Figure 2.**
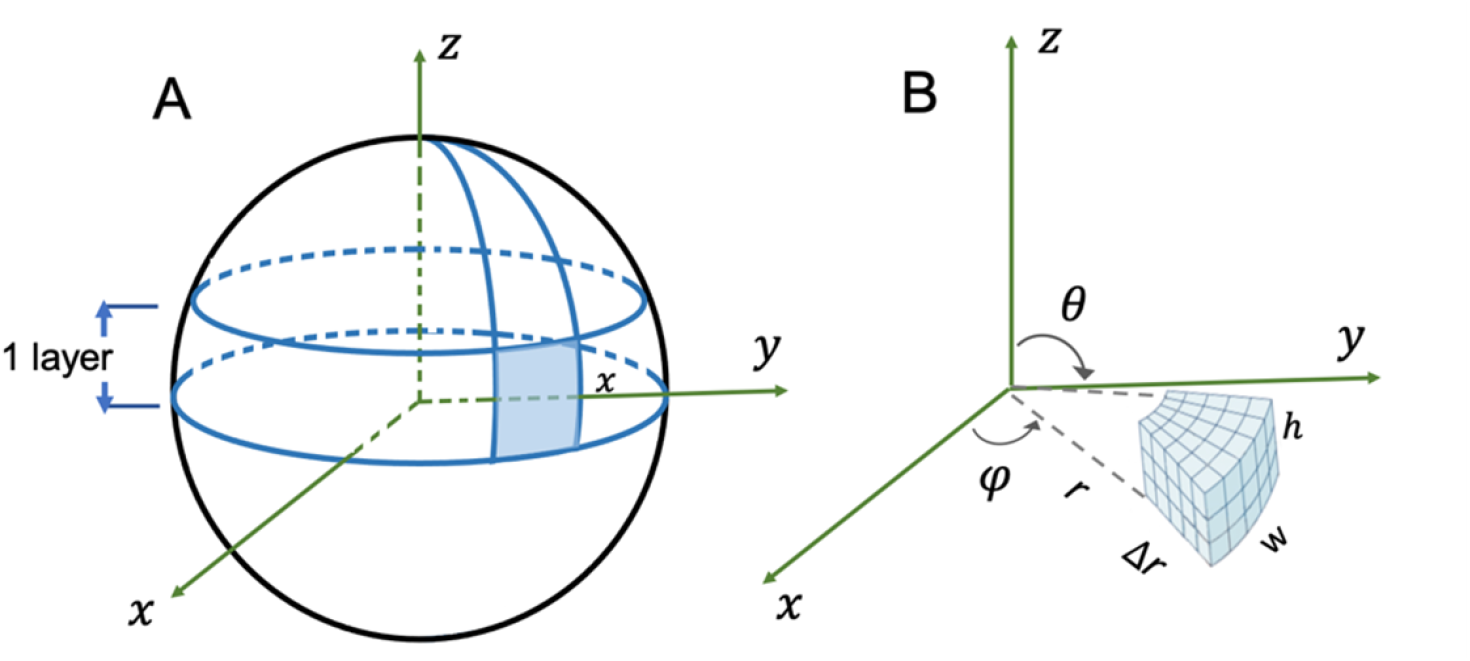
IIllustration of magnetic blocks. Magnetic materials were prescribed only on top half of the sphere. There was a total of 7 layers. Each layer was subdivided into blocks. Grey area in (A) is expanded in (B) to illustrate one block. The program discretizes each block into small sections (meshes) for electromagnetic calculation.

The magnetic blocks were generated using the CoilCore functions in the BEM-FMM package. First, a polygon that depicted the base shape of the magnetic block was defined and meshes were generated; second, the meshed plane was extruded upward to the height *h* (see figure 2). Since each discrete block is vertical to its base, layer 6 leaves a circular opening at the top. So an additional layer (layer 7), a circular block (“top cap”) to cover this opening, was added such that the entire top hemisphere was surrounded by discrete magnetic blocks.

Thus, the total number of discrete magnetic block was 65; layers 1 to 6 had 64 blocks, symmetrically distributed in the *X*–*Y* plane, with one additional layer (layer 7) at the top. Equation 5 mathematically reduces to finding the optimum thickness for each magnetic block (Δ*r*_*i*_, *i* = 1 … 65), such that the cost function is minimized.

### 2.6. Cost function definition

The region of interest (ROI) was prescribed as a cone inside the spherical head model (see figure 1) defined by the equation

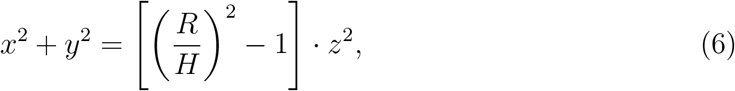

where *R* = 70 *mm* and *H* = 60 *mm*. *H* and *R* together define the 3D surface of a cone. *z* ranged from 30 to 60 *mm* with a step size of 2 *mm*. The points of interest on the circle corresponding to a given *z*_*j*_ was calculated with a step size in *x* coordinates of 3 *mm*, resulting in a total of 166 points. The ideal E-field was defined as a linear decay along the cone surface (i.e. along the radial direction toward the center of the sphere). While the E-field generally decays superalinearly, setting the ideal E-field as a linear decay moves the optimization towards this goal (see below).

### 2.7. Optimization procedures

Optimization of the thickness of each block was done using Newton’s gradient descent method. The procedure was as follows: the variables for each magnetic block were parametrized, and the E-field calculation was looped through each magnetic piece. The procedure was then repeated with 5 *mm* added to each block’s thickness. The gradient, or each piece’s contribution to the cost function, noted as *G_i_*, was calculated by computing first partial derivative of the error function over each block (see equation 5). The thickness of each individual block was then adjusted based on its gradient. The above processes were repeated 3 times to achieve the final results. All the computation was performed using MATLAB (MathWorks).

### 2.8. Experimental validation

As a proof-of concept, we mapped the E-field generated by a double-cone coil with and without magnetic materials. A clear acrylic globe (16 *cm* in diameter) was used as the head model; iron blocks with a purity 99.98% was used as the magnetic material. High purity iron has high relative permeability (1000–2000) and was machined to fit the spherical surface of the globe. Figure 3 shows the measurement setup. The coil and the stimulator (Model: Magstim 200) were both from Magstim Inc. U.K. The globe was filled with saline water by dissolving sodium chloride into water at 0.9% concentration to mimic the conductivity of human tissue [22].

**Figure 3.**
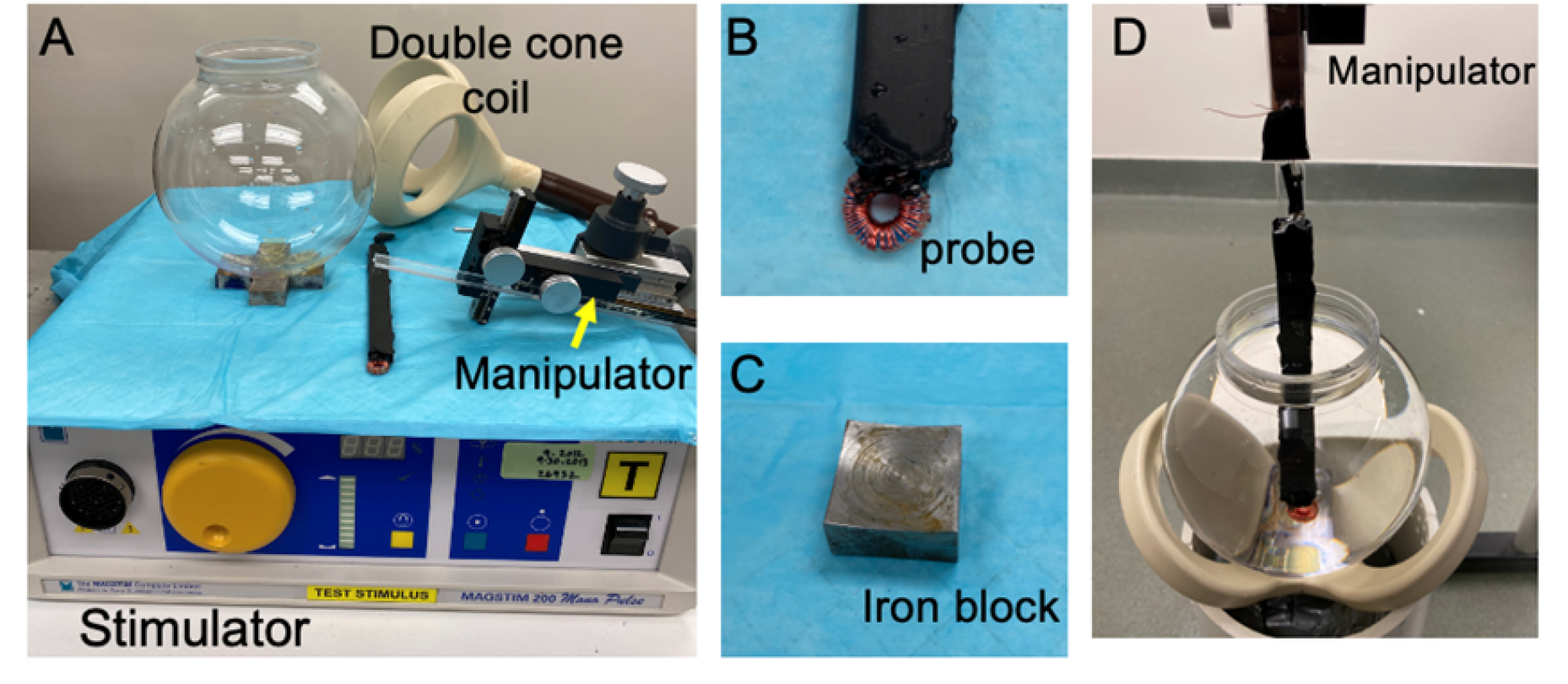
E-field measurement setup. (A): a commercial double cone coil driven by a stimulator was used to deliver the TMS pulse; a manipulator with 0.1mm precision was used to move the probe along *X*, *Y*, and *Z*. Measurements were made with and without the iron blocks while keeping the space between the coil and head phantom the same. (B): a homemade probe (Rogowski coil) was used to measure the E-field. (C): high purity iron (99.98%) block was machined to fit the spherical surface of the head phantom. (D): the globe filled with water containing 0.9% sodium chloride. The probe and coil were in position for measurement.

An E-field probe (Rogowski coil) was handmade to detect the E-field (figure 3B) [23]. The coil was carefully wound such that it canceled out the magnetic field signal, being sensitive only to E-field induced by the magnetic field. A toroid core was used to enhance the sensitivity of the probe and an oscilloscope was used to read the probe’s signal. E-field was measured with and without the iron blocks. Special care was taken to keep all other parameters the same. The iron blocks were fixed on the bottom of the globe using hot gun glue.

Based on theoretical studies (see results), it was found only the blocks close to top of the model (near the coil) had a strong effect on the E-field. Since the purpose of this experiment was only to validate the principle of my coil design strategy, only 5 blocks were used in this preliminary study.

## 3. Results

### 3.1. Cross-validation of E-field calculation with known winding patterns in commercial TMS coils

To compare our E-field calculations with the BEM-FMM software package and our quantifications of coil penetration depth (*d*_1/2_) and field spread (*s*_1/2_), four widely used commercial coils were selected from Magstim Co. Ltd, UK. These coils have known design parameters [10, 16], which were modeled in the BEM-FMM software package. Results are summarized in table 1, labled as “Present” values. For comparison, literature values calculated using finite-element simulation software package are also listed [16]. In general, both the *d*_1/2_ and *s*_1/2_ values are in good agreement with literature data with a difference of 2.02%±0.98% in *d*_1/2_ and 3.12%±3.58% in *s*_1/2_ (mean±standard deviation).

**Table 1.**
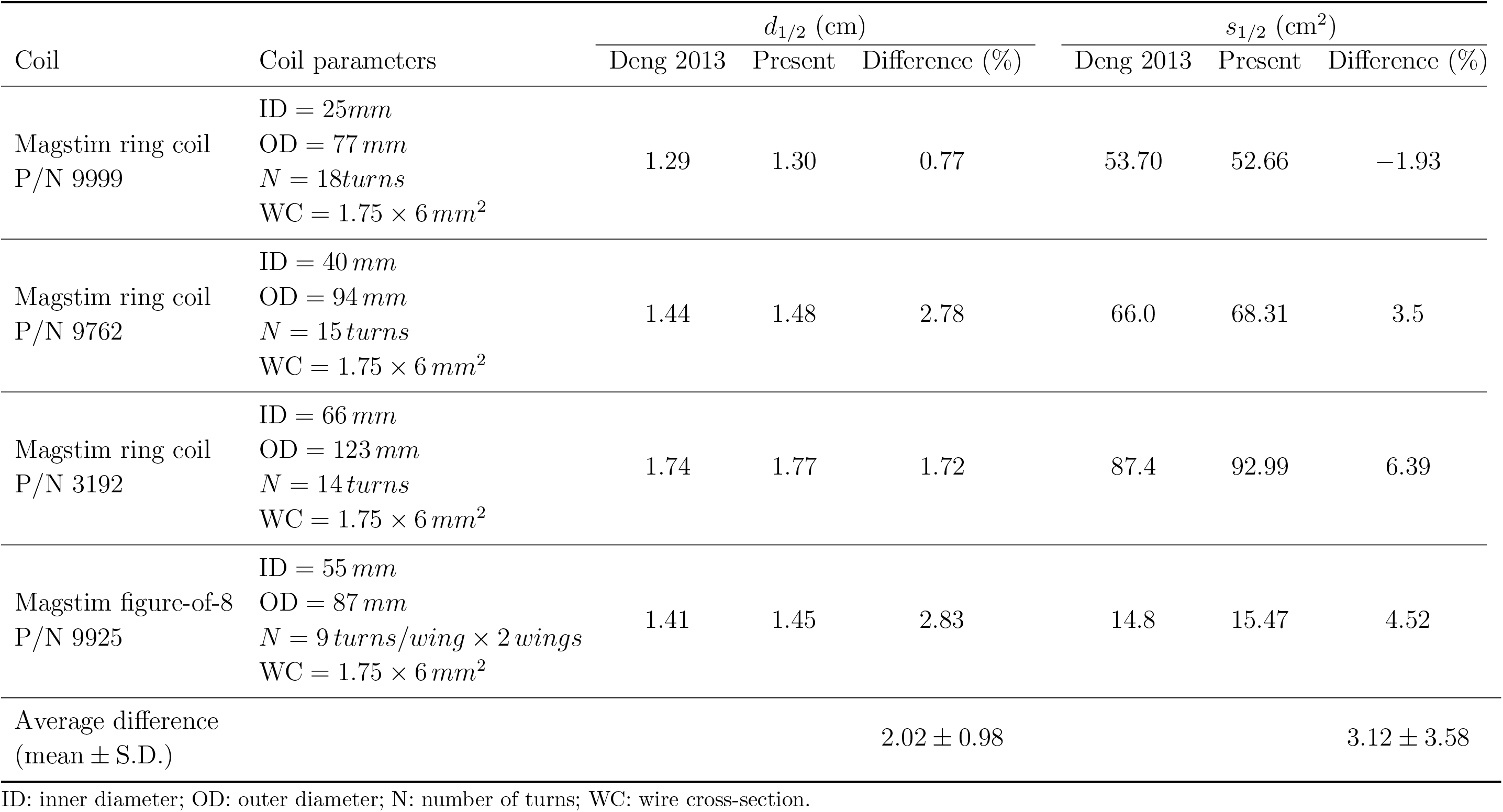
Comparisons of E-field penetration depth (*d*_1/2_) and spread (*s*_1/2_) with literature data

### 3.2. Coil optimization

We calculated the E-field by the double-cone coil in the presence of each magnetic block. As one example, figure 4 shows the E-field distribution in the presence of one block. The magnetic material clearly changed the E-field as indicated by the white arrow, which would otherwise be symmetric in the absence of the magnetic block.

**Figure 4.**
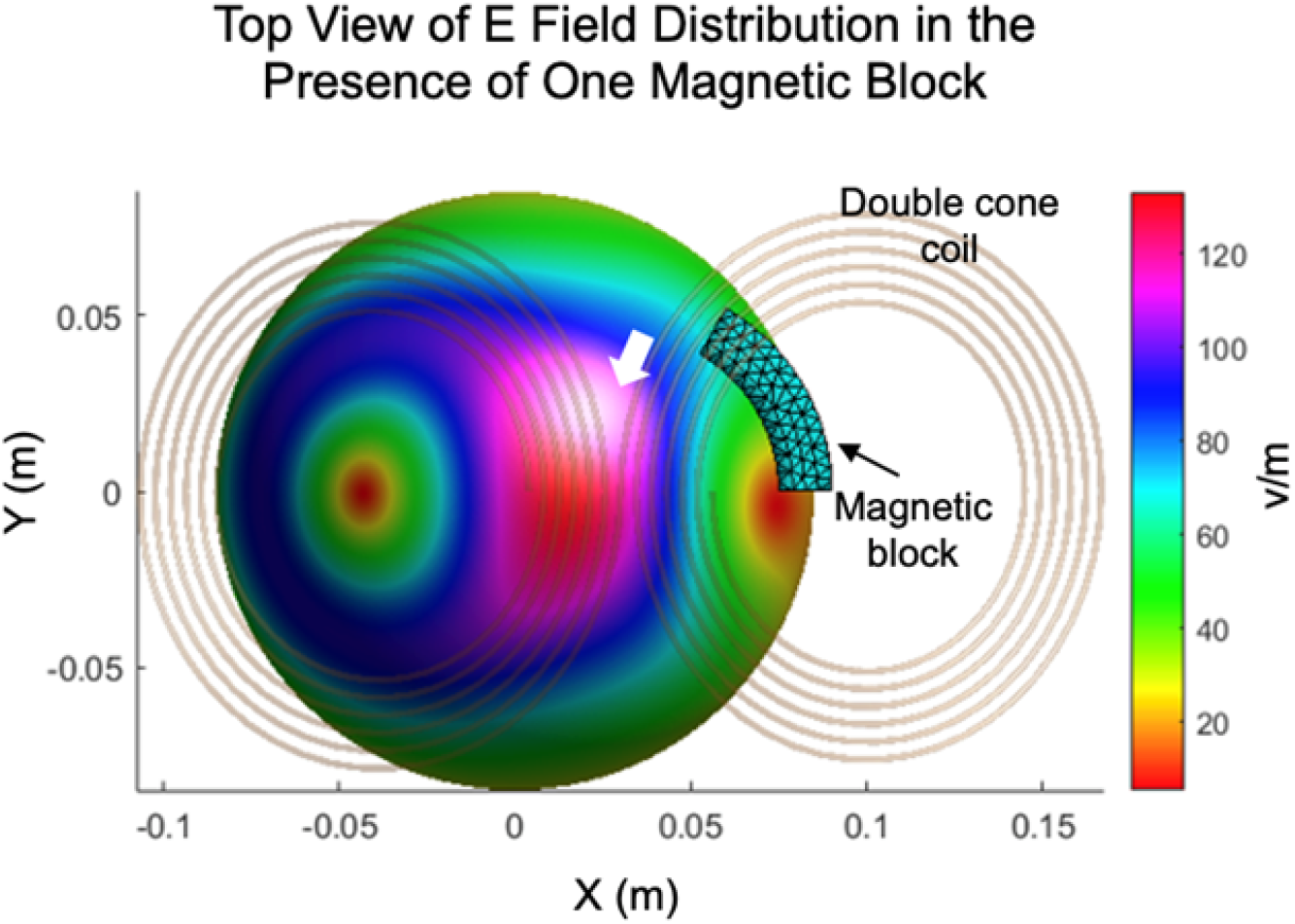
Illustration of E-field calculation in the presence of a magnetic block. Double-cone coil generates symmetric field in the spherical head model (indicated by the green ball). White arrow indicates field perturbation by the magnetic block, which breaks the symmetric distribution of the E-field generated by the double-cone coil.

To quantify the contributions of the 65 blocks to the cost function, the thickness of each block was increased by 5 *mm*, i.e. d*r* = 5 *mm*; and repeated the above computation process one more iteration. The gradient, which was the difference in cost function caused divided by d*r* was then computed in MATLAB. Figure 5 shows gradients of individual blocks. Note that since blocks from layers 1 to 6 are symmetric in the *X*–*Y* plane, the gradients from only half of these blocks (*n* = 32) and the block on layer 7 (“top cap”) are shown.

**Figure 5.**
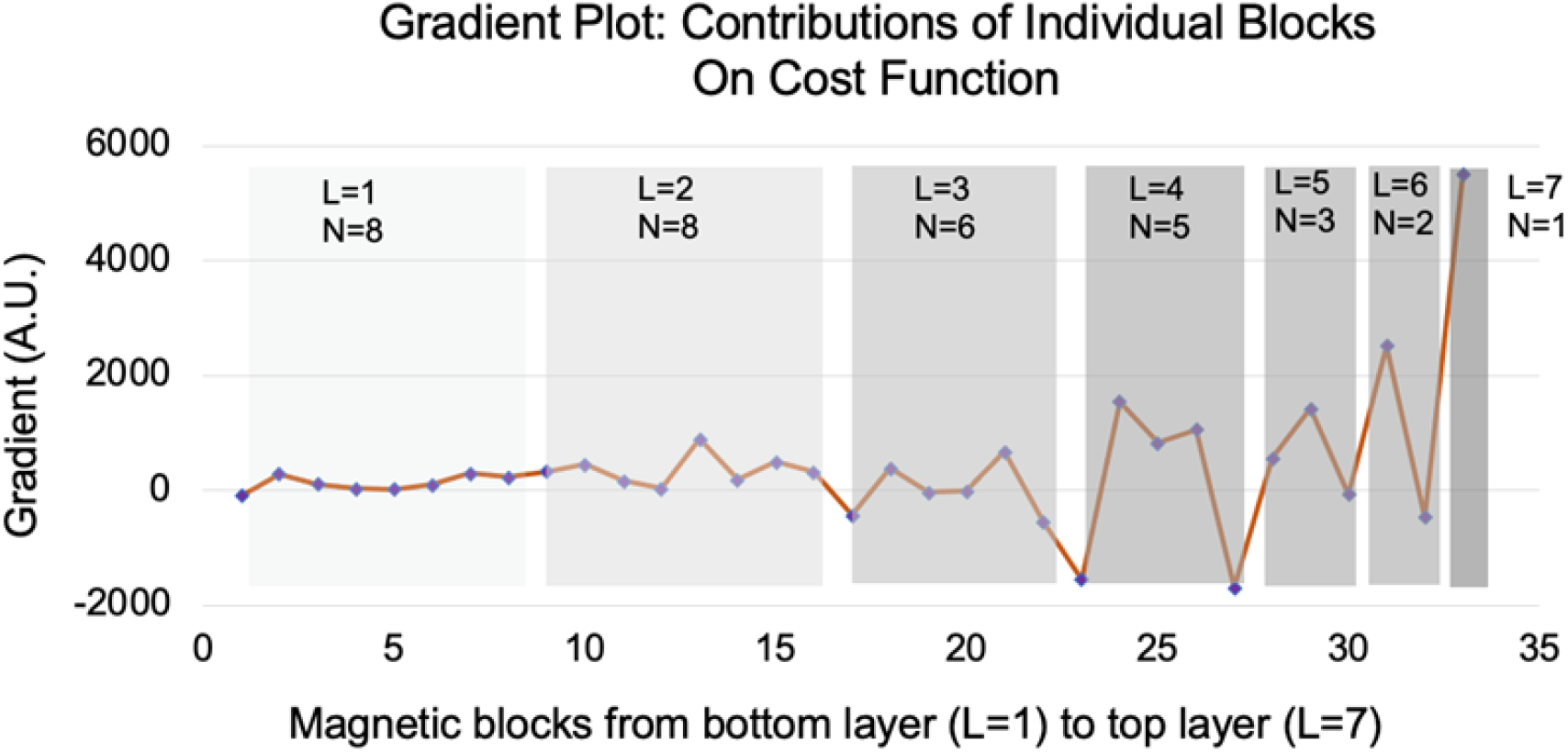
Effects of magnetic blocks from bottom layer (L1) to top layer (L7) on the cost function. Blocks from layers 1 to 6 are symmetric in the *X*–*Y* plane, only half of these blocks (*n* = 32) plus 1 block in L7 are plotted. Magnetic blocks at layers 4–7 had a greater contribution than those at layers 1 to 3. The gradient plot is in arbitrary unit (A.U.).

Figure 5 shows that blocks in layers 4–7 had higher gradients than in layers 1–3, which was expected, since the latter ones are farther away from the coil. Of a particular note, layer 7, the “top cap”, had the greatest contribution to the cost function.

To further investigate the effects of the “top cap” block on the E-field, figure 6 shows E-field along the center line with and without this block. This E-field decayed exponentially in the absence of the “top cap” (red curve). The blur curve (with “top cap”) decayed slower, but its peak E-field strength was reduced by about 55%.

**Figure 6.**
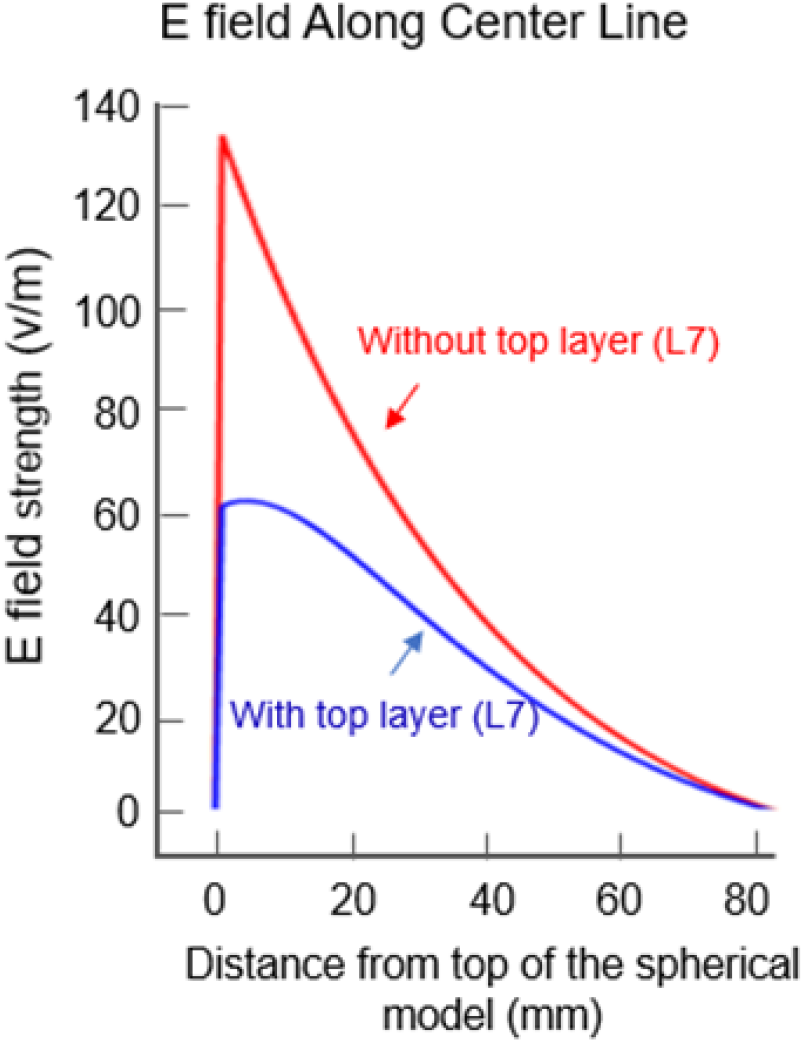
E-field plots along the center line of the spherical head model. The magnetic block at the top layer (layer 7, “top cap”) had a strong effect on both the decay rate (penetration depth) and the strength of the E-field.

Thus the “top cap” clearly slowed the E-field decay rate, leading to greater penetration depth at the cost of E field strength. Using Newton’s gradient decent method, the thickness of individual blocks was adjusted based on their gradients and repeated the above computation steps for 3 iterations. Both *d*_1/2_ and *s*_1/2_ were computed at the end of each iteration. Table 2 summarizes the final results. For comparison, data from commercial coils that have been designed specifically for stimulating deep structures are also listed. Our design achieved a better or comparable penetration depth (*d*_1/2_), with much smaller field spread (*s*_1/2_), as revealed by the ratio of *s*_1/2_ over *d*_1/2_ in Table 2 the current design has a ratio of 23.35, while the Brainsway H1 coil has a ratio of 51.61, and Magstim cap coil has a ratio of 63.38. Thus our design improves coil depth–focality tradeoff by a factor of 2.2 to 2.7 better than these commercial coils.

**Table 2.** Comparisons with coil depth–focality tradeoff

|  | Current design | Commercial deep coils<sup>1</sup> |  |  |
| --- | --- | --- | --- | --- |
|  |  | Brainsway H1 | Brainsway H2 | Magstim cap |
| $d_{1/2}$ | 2.60 | 2.17 | 2.23 | 2.63 |
| $s_{1/2}$ | 60.72 | 112 | 131 | 168 |
| Tradeoff ($s_{1/2}/d_{1/2}$) | 23.35 | 51.61 | 58.74 | 63.88 |
| Tradeoff improvement | 1 | 2.2 | 2.5 | 2.7 |

<sup>1</sup> Data are from Deng et al. 2013 [16]. These coils are specifically designed for stimulating deep brain regions: H1 and H2 coils are from Brainsway, Israel; Cap coil is from Magstim Co. Ltd, UK. Tradeoff improvement is the Tradeoff value of a commercial coil relative to the current design.

### 3.3. Experimental validation

As a proof-of-concept, E-field inside a spherical phantom filled with saline water was measured. Figure 7A shows a typical E-field waveform using the homemade E-field probe. There were brief artifacts at pulse initiation, lasting for about 4 μ*s*, which was caused by the stimulator system. The first peak following the artifacts was caused by sharp increase in coil current and was used to indicate E-field strength. Figure 7B shows E-field along the center line of the spherical phantom at 5 *mm* steps. The E-field decay curve in the absence of iron blocks can be well fitted with the 4^th^ order polynomial *y* = 0.00006*x*^4^ − 0.0173*x*^3^ + 1.8452*x*^2^ − 96.234*x* + 2298.8. (*R*^2^ = 0.9988). The E-field in the presence of iron blocks can be fitted with the 6^th^ order polynomial *y* = −0.0000002*x*^6^ + 0.00005*x*^5^ − 0.0061*x*^4^ + 0.3387 − 9.2053 + 85.418*x* + 854.98. (*R*^2^ = 0.9985). Also note that despite the small factors in the 4^th^, 5^th^ and 6^th^ orders of the polynomials, these factors were significant since the power operations of *x* in the range of 10–85 (*x*-axis) resulted in significant values. The blue and red arrows in Figure 7B indicates *x* locations when the E-field strength decayed by 50%.

**Figure 7.**
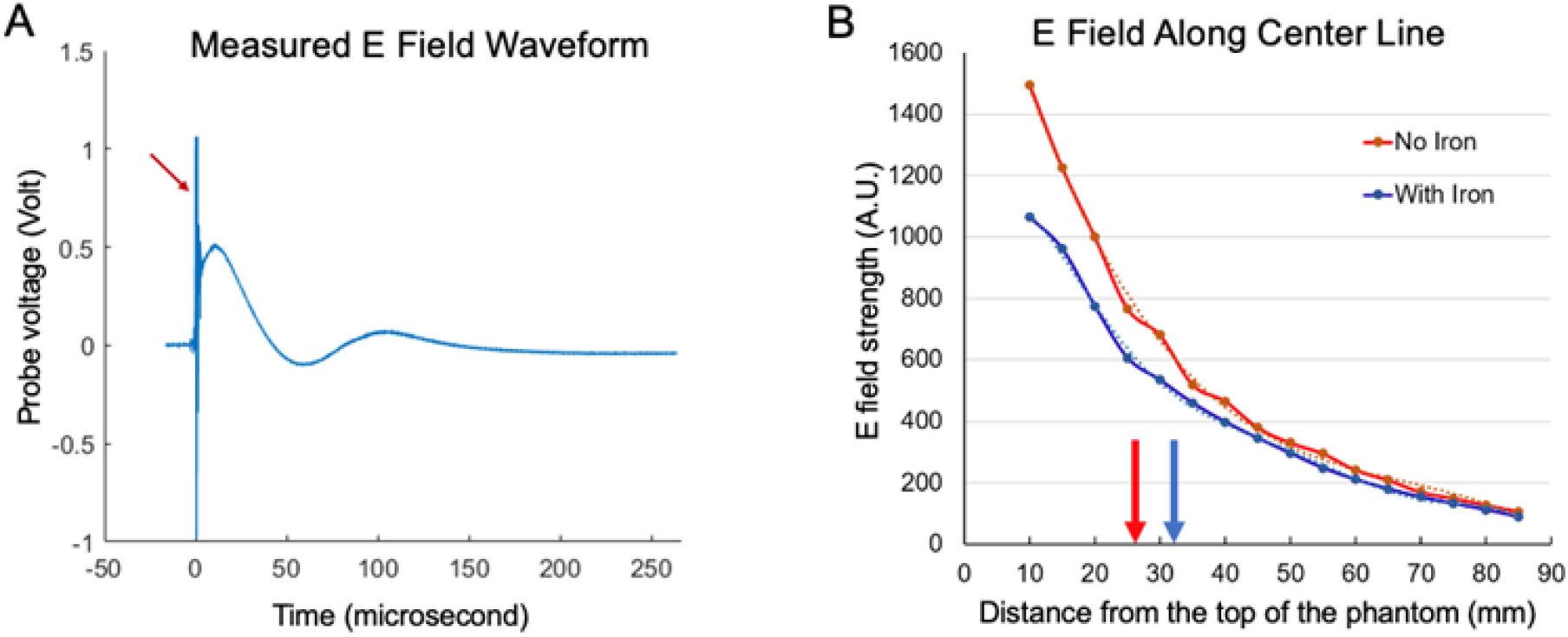
E-field measurement in a phantom filled with saline. A) An exemplar of the E-field waveform measured with the customized probe. The red arrow in (A) indicates artifacts caused by the stimulator system when pulse started. B. Measured E-field distribution along the center line of a spherical head phantom. The dash lines are corresponding trendlines fitted with polynomials. The red curve (no iron blocks) could be well fitted with a 4^th^ order polynomial; the blue curve (with iron) could be well fitted with a 6^th^ order polynomial. The red and blue arrows in (B) indicates *X* location when the field dropped to 50% of the peak values (at *x* = 10), respectively. Iron blocks influenced both the decay rate (penetration depth) and the strength of the E-field.

The E-field decay curves suggest that iron blocks favorably slowed decay rates, leading to improved penetration depth. Also shown in figure 7B is that iron blocks reduced peak E-field strength by 29% (at *x* = 10 *mm*). These results are largely in agreement with theoretical data shown in figure 6.

## 4. Discussion

In the present study, a novel coil augmentation strategy is proposed, which applied magnetic materials to shape the electromagnetic field generated by conventional coils. Through numerical optimization, we achieved coil penetration depth better than state-of-the-art deep coils while enhancing the coil focality, improving the depth–focality tradeoff by a factor of 2.2 to 2.7 (table 2).

As a proof-of-concept, E-field inside a spherical head phantom was mapped with and without magnetic materials. The experimental results confirmed that the use of magnetic materials reduced E-field decay rate, consistent with theoretical calculation (figures 6 and 7).

### 4.1. Limitations

Due to technical difficulties in machining magnetic materials, only 5 blocks of equal thickness were used in this experiment. This was based on the observation that only the blocks close to top of the model had strong effect on the E-field (figure 5), and the purpose of this experiment was to validate the principle of the new coil design strategy. Further improvement in coil performance can be expected if the shapes of the iron blocks follow the design patterns. Nevertheless, the experimental results strongly support the theoretical analysis.

The improvement in penetration depth–focality tradeoff was at the cost of reduced E-field strength by about 29%. This means that to reach the same E-field strength, a stronger current is necessary, which will lead to more coil heat. As a result, more efficient coil cooling is warranted. Weak surface E-field has advantages, however. Davey and Riehl [24] suggested that weakening surface E-field could reduce TMS-induced pain sensation in patients, and thus is preferable for clinical applications.

In summary, magnetic materials are proposed to optimally shape the electric field produced by conventional TMS coils. Theoretical analyses and experimental data demonstrate that this approach can substantially improve coil penetration depth– focality tradeoff—a major technical limitation in existing TMS system. This novel coil design has the potential to improve clinical outcome of TMS treatment.

## Acknowledgments

CL would like to thank his research teacher Ms. Ireland for her guidance. ZD was supported by the National Institute of Mental Health Intramural Research Program (ZIAMH002955).

